# Ggtree: a serialized data object for visualization of phylogenetic tree and annotation data

**DOI:** 10.1101/2020.10.21.348169

**Authors:** Guangchuang Yu

## Abstract

Ggtree supports mapping and visualizing associated external data on phylogeny with two general methods. The output of ggtree is a ggtree graphic object that can be rendered as a static image. Most importantly, the input tree and associated data that used in visualization can be extracted from the graphic object, making it an ideal data structure for publishing tree (image, tree and data in one single object) and thus enhance data reuse and analytic reproducibility.

---

Phylogenetic data have enormous potential for reuse as phylogeny is becoming central to a wide range of research in ecology, evolutionary biology, epidemiology and molecular biology. Reusing phylogenetic data can contribute to synthesize phylogenetic knowledge and comparative analyses in a number of scientific disciplines. However, a previous survey alerts that ~60% of published phylogenetic data are lost to science forever^1^. One of the reason for this situation is that phylogenetic trees are often published as static images and lack of interoperable file format for data sharing^2^. Creating tree figures annotated with associated data (taxonomy information, meta-data, phenotypic and epidemiological data, etc.) is a routine practice. Although tools for tree visualization and annotation are proliferating, the dominant objective remains to produce a publication-ready figure, which involves multiple steps in selecting the annotation data (*e.g.*, bootstrap values) and rendering it on the tree (*e.g.*, as text labels or branch colors). The process is one-way and a dead end to yield a static figure that the underlying information cannot be reused. We need a paradigm shift from producing a static figure to a serialized data object that contains the tree, associated data and visualization directives in addition to render as a visualization graphic.

Here, we describe a data structure, the *ggtree* object, defined in the *ggtree* package. *Ggtree* is an R/Bioconductor package for the visualization and annotation of phylogenetic tree with diverse associated data^3^. We previously proposed two methods for mapping and visualizing associated data on phylogeny^4^. Taxon species associated data can be linked to the tree structure within the *ggtree* object by the *%<+%* operator and complex associated data can be visualized by specific *geom* layer in separate panel and align to the tree based on the tree structure using *facet_plot* function^4^. Data that mapped and visualized using these two methods are preserved and can be extracted from the *ggtree* object. In Fig. 1, an associated data was attached to the tree using the *%<+%* operator and the posterior values were used to color circle points on the tree (Fig. 1A). The output *ggtree* object is a graphic object and can be rendered as a static figure (Fig. 1B). The object preserves information of the phylogenetic tree as well as associated data that was attached (Fig. 1C). Users can convert this graphic object to a *phylo* (only tree structure without annotation data) or *treedata* (tree structure and annotation data) object. The tree object can be further processed using *tidytree* or *treeio* packages and can be exported to Newick, Nexus or BEAST Nexus which enables associated data to be stored as annotated elements^5^. The *ggtree* graphic object also contains visualization directive and the visualization style that can be reused to visualize a new tree object (Fig. 1D), which is similar to Microsoft Word Format Painter. Data that used in *facet_plot* can be complex and heterogeneous, such as genetic information at pan-genome scale and species abundance distributions. Although the data is not directly mapped to the tree, it can also be extracted from the *ggtree* object. The body weight information was used to be visualized as barplot with the tree side by side (Fig. 1E) and it can be extracted from the *ggtree* graphic object using *facet_data* function (Fig. 1F).

**Fig. 1.**
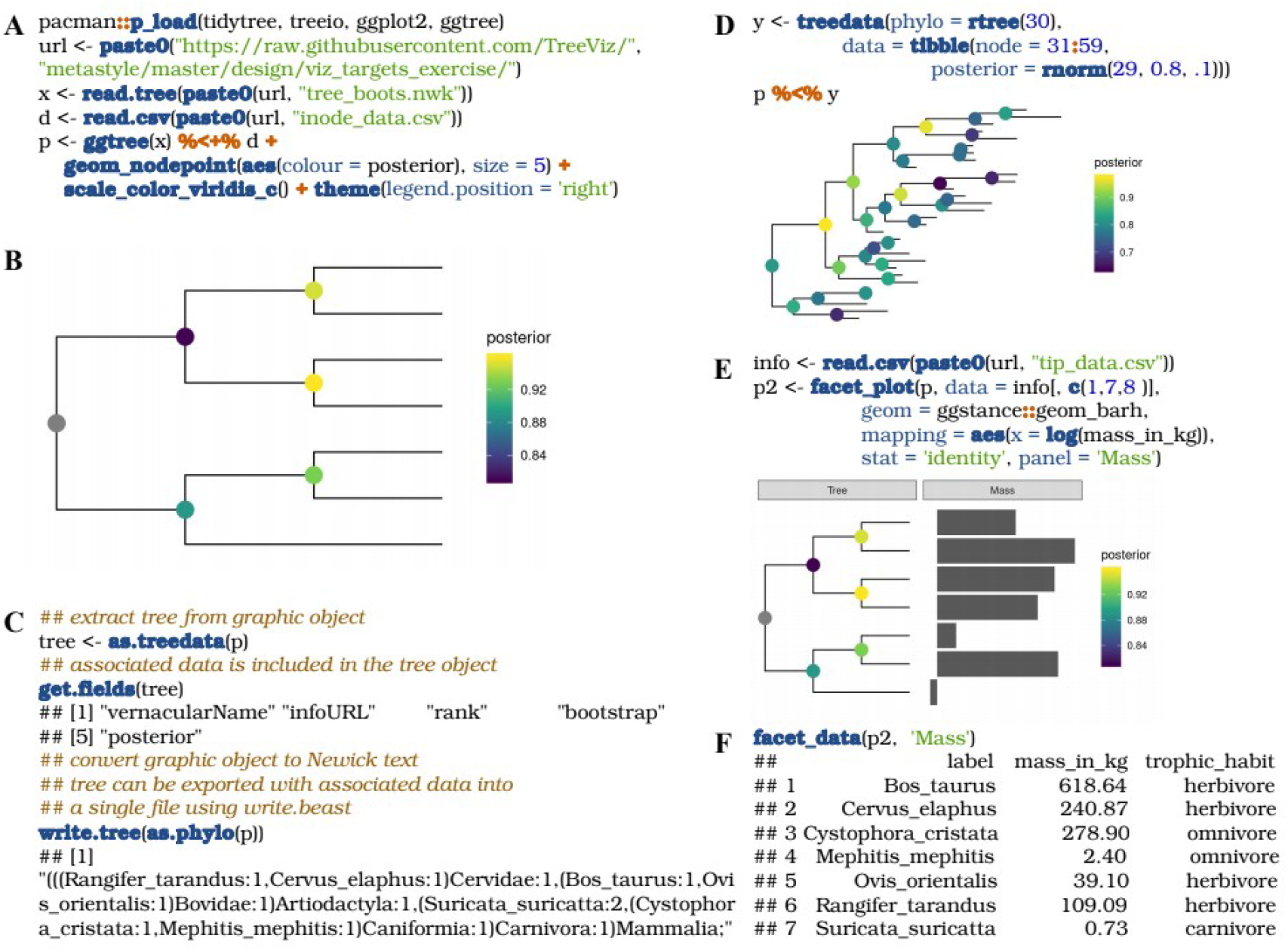
Examples of using *ggtree* object to store phylogenetic tree, associated data and visualization directive. A tree in Newick format was read into R as *x* and an associated data, *d*, that stores posterior of internal nodes were used to visualize circle points that colored by the posterior values. The output graphic object, *i.e.* the *ggtree* object, was stored in *p* (A). The *ggtree* object, *p*, can be rendered as a picture by *print(p)* (B). The *ggtree* object contains the information of phylogenetic tree and associated data. It can be converted back to a *treedata* object with the tree and all the associated data, including posterior, bootstrap and other values in *d* that were attached to the graphic object, *p*. The tree can be exported to Newick text (without associated data) and BEAST Nexus file (with associated data) (C). The *ggtree* object contains visualization directive which can be reused to visualize a new tree object by applying its visualization style (D). Associated data can be visualized with the tree side by side using *facet_plot* (E). The data that used in *facet_plot* can also be extracted from the graphic object using *facet_data* function (F).

In addition to the tree itself, all the information that are mapped and visualized on the phylogeny using the methods proposed in *ggtree*^4^ are reusable, including data that rendered as visual characteristics to display the tree and data that used to produce graph to align with the tree. With *ggtree*, phylogenetic data as well as the visual directives that used to create the image in publication are seamlessly stored in a single object. Making the visualization more portable and transparent and ensuring high quality phylogenetic information to be shared and reused in different projects. It also offers great potential for remote collaborators to modify tree data presentation. If a tree was published using the *ggtree* graphic object (*i.e.*, as supplemental file or deposited to data repository), others can download and read the file into R to reproduce the figure that exactly identical to the published one and are able to add or modify layers of tree annotation by reusing data in the object or from other sources. Users can obtain tree structure and associated data from the *ggtree* object for integrative and comparative analyses using a number of R packages. A paradigm shift from generating static figure to producing serialized data object like *ggtree* not only enhances analytic reproducibility but also assures data reusability and facilitates synthetic and comparative studies to identify and pursue new questions. As phylogenetic data from various discipline are becoming widely available, this paradigm shift should be a criteria for designing next-generation tree visualization tools.

## Code availability

*ggtree* is freely available at https://www.bioconductor.org/packages/ggtree. A complete reference of *ggtree* with many step-by-step examples on how to process and visualize phylogenetic data is available in the online book, https://yulab-smu.top/treedata-book/.

## Acknowledgement

This work was supported by Startup fund from Southern Medical University (G618289088).

## References

Magee A.F. et al. PLOS ONE. 9:e110268. (2014)

Cranston K. et al. PLoS Curr. [Internet] 6. Available from: https://www.ncbi.nlm.nih.gov/pmc/articles/PMC4073804/ (2014)

Yu G. et al. Methods Ecol. Evol. 8:28–36. (2017)

Yu G. et al. Mol. Biol. Evol. 35:3041–3043. (2018)

Wang L.G. et al. Mol. Biol. Evol. 37:599–603. (2020)

